# Bacillus anthracis chain length, a virulence determinant, is regulated by a transmembrane Ser/Thr protein kinase PrkC

**DOI:** 10.1101/2020.03.15.992834

**Authors:** Neha Dhasmana, Nishant Kumar, Aakriti Gangwal, Chetkar Chandra Keshavam, Lalit K. Singh, Nitika Sangwan, Payal Nashier, Sagarika Biswas, Andrei P. Pomerantsev, Stephen H Leppla, Yogendra Singh, Meetu Gupta

**Author notes:** These authors contributed equally to this work (Neha Dhasmana, Nishant Kumar, Aakriti Gangwal, Chetkar Chandra Keshavam, Lalit K. Singh).

## Abstract

Anthrax is a zoonotic disease caused by *Bacillus anthracis*, a spore-forming pathogen that displays a chaining phenotype. It has been reported that in a mouse infection model, systemic inoculation with longer bacterial chains caused blockade in lung capillaries. The blockade resulted in increased pathophysiological consequences viz, hypoxia and lung tissue injury. Hence, chaining acts as a virulence factor and molecules that regulate the chaining phenotype can be the potential drug targets. In this study, we have identified the serine/threonine protein kinase of *B. anthracis*, PrkC, localized at the bacteria-host interface, as a determinant of bacterial chain length. *In vitro, prkC* disruption strain (BAS Δ*prkC*) grew as shorter chains throughout the bacterial growth cycle as observed through phase-contrast and scanning electron microscopy. Since molecules such as BslO, a septal murein hydrolase, that catalyzes daughter cell separation and Sap, an S-layer structural protein required for the septal localization of BslO, are known to influence chain length, a comparative analysis to determine their levels was done through western-blot analysis. Both BslO and Sap were found to be upregulated in BAS Δ*prkC* at the majority of the time points. Additionally, PrkC disruption was observed to have a significant effect on bacterial growth and cell wall thickness. In BAS Δ*prkC* strain, a decrease in the cell wall thickness and an increase in the multi-septa formation was observed through transmission electron and confocal microscopy respectively. Altogether, we show that PrkC disruption affected chaining phenotype, cell growth and cell wall thickness and also report that the associated molecules were de-regulated. Through this work, we show for the first time that the chaining phenotype is regulated by PrkC, a transmembrane kinase with a sensor domain. During infection, PrkC may regulate the chaining phenotype through the identified signaling mechanism.

**Authors summary:** *B. anthracis*, a spore-forming pathogen is the causative agent of anthrax, a zoonotic disease that primarily affects livestock and wildlife. Humans are at risk of contracting this disease through exposure to spores generated by infected animals. In the past, *B. anthracis* spores have been used as a bioterror agent. Hence, there has been a continuous effort to understand the biology of this pathogen to develop both therapeutic and prophylactic treatment. Various virulence factors that are essential for *B. anthracis* pathogenesis have been identified. The ability of *B. anthracis* to grow in chains acts as a virulence factor. Longer bacterial chains are reported to cause blockade of lung capillaries in the mouse infection model. In this study, we have shown that the disruption of the lone serine/threonine protein kinase, PrkC, localized at the bacteria-host interface leads to the shortening of the bacterial chains. We have seen that the depletion of PrkC results in an increase in the levels of the proteins responsible for de-chaining. Also, we have analyzed the effect of the disruption on cell growth, bacterial cell wall and septa formation. Since PrkC is a surface localized kinase with an extracellular domain that lacks homology to human proteins, it can be a target for new drugs. Disruption of PrkC activity and hence the longer chains *in vivo* may prevent pathophysiological consequences associated with the capillary blockade.

## Introduction

Bacteria exhibit diverse shapes and morphologies, the result of long evolutionary processes that select genotypes best suited for bacterial survival. Apart from the variation in shape, bacteria display multi-cellular structures such as aggregates, biofilms, and chains/filaments [1, 2]. *Bacillus anthracis*, the Gram-positive spore-forming pathogen of grazing mammals, and the etiological agent of anthrax grows as chains of rod-shaped cells [3-6]. In the environment, *B. anthracis* persists primarily as metabolically inert oblong spores. Germination happens in the presence of an optimal signal within the host [7]. Emerging evidence indicates that spores can germinate, multiply and persist even outside their vertebrate host, in the presence of a nutrient-rich environment in the root rhizosphere and simpler biological systems such as earthworm, housefly, and amoebae [6, 8-11].

*B. anthracis* spores can infect humans through three routes – gastrointestinal, inhalational, and cutaneous [7, 12-14]. The highest mortality rates are seen in inhalational anthrax [15]. Multiple factors act as virulence determinants [16-18]. However, the secreted binary exotoxins (lethal toxin and edema toxin) and the anti-phagocytic poly-γ-D-glutamic acid capsule, encoded by the virulence plasmids pXO1 and pXO2, respectively, act as the primary virulence factors [7, 19, 20]. The exotoxins perturb host immune responses and cause toxemia, and the capsule prevents engulfment by phagocytes, resulting in septicemia [20, 21].

Among other factors that play a role in virulence, the bacterial chaining phenotype has been shown to contribute significantly [4, 5]. During initial stages of infection, *B. anthracis* spores phagocytosed by macrophages, germinate, and grow in chains before causing cell rupture [22]. In mice, the high pathogenicity of systemically inoculated *B. anthracis* strain making capsule but not the toxins (encapsulated but nontoxinogenic strain) was linked to chain length-dependent blockade of alveolar capillaries leading to hypoxia, lung tissue injury, and death [4, 5]. Of note, the lung is the terminal organ targeted by *B. anthracis*, irrespective of the route of infection [5, 23, 24]. These studies indicate that the chaining phenotype presents a survival advantage to *B. anthracis* within its host during both early and late stages of infection.

Intrigued by the relevance of this morphotype in the biology of *Bacillus* species, various groups have tried to identify the mechanisms controlling bacterial chain length in both pathogenic and non-pathogenic strains [25-33]. In *B. anthracis*, one of the determinants of bacterial chain length is the septal peptidoglycan hydrolase, BslO (*bacillus s*urface *l*ayer *O*). BslO is a *Bacillus S*-*l*ayer associated protein (BSL) with *N*-acetylglucosaminidase activity that catalyzes daughter cell separation [28]. Restrictive deposition of BslO to the septal region is, in turn, attributed to sequential coverage of the cell wall by the primary S-layer proteins (SLPs), Sap (*s*urface *a*rray *p*rotein) and EA1 (*e*xtractable *a*ntigen *1*) [27]. SLPs and BSLs associate with the pyruvylated secondary cell wall polysaccharides (SCWP) through their conserved S-layer homology domain (SLH) [34, 35]. While several enzymes that influence the chaining phenotype through their role in synthesis/modification of SCWPs and hence the attachment of SLPs to SCWPs have been identified in *B. anthracis* [35, 36], a sensory molecule with a potential to regulate the chaining phenotype, possibly through regulation of one of these factors, remains unknown.

Through this work, we identify *B. anthracis* PrkC, the only serine/threonine protein kinase (STPK) localized at the bacteria-host interface, as a determinant of bacterial chain length. We show that the *B. anthracis* Sterne 34F2 *prkC* mutant strain (BAS Δ*prkC*) is not able to attain a chaining phenotype throughout the bacterial growth cycle. Both BslO and Sap are found to be upregulated in BAS Δ*prkC*, which probably creates a condition that favors de-chaining. Additionally, PrkC is also shown to influence the bacterial cell division, possibly through the regulation of the cytoskeletal protein, FtsZ. Through this work, we propose that PrkC, a transmembrane kinase with a sensor domain, perceives growth permissive signals and maintains the levels of the primary proteins involved in de-chaining to regulate the chaining phenotype.

## Results

STPKs, earlier thought to be limited to eukaryotes, are now identified as integral components of the bacterial systems with a definite role in bacterial survival and pathogenesis [37]. In *B. anthracis*, three STPKs have been characterized, namely PrkC (BAS 3713), PrkD (BAS 2152), and PrkG (BAS 2037) [38-40]. Among these, PrkC is the only membrane-associated protein kinase [39, 41]. The extracellular ligand-binding motif of PrkC is composed of peptidoglycan binding PASTA (*p*enicillin-binding proteins *a*nd *S*er/Thr kinase-*a*ssociated) repeats. Interaction of the PASTA domain with peptidoglycan was first demonstrated for PrkC, wherein it was shown to interact with peptidoglycan fragments generated by neighboring growing cells, thereby triggering germination of *B. subtilis* and *B. anthracis* spores [42]. Apart from a role in germination, *B. subtilis* PrkC has been implicated in stationary phase processes, cell wall metabolism, cell division, sporulation, germination, and biofilm formation [41, 43-47].

### *prkC* disruption results in bacteria with shorter chain length

Previously, our group had shown that *B. anthracis* PrkC-mediated processes play an essential role in germination and biofilm formation [48, 49]. Some of the components of the PrkC-mediated signaling cascade leading to these processes were identified [48. 49]. Work done by other groups implicated *B. subtilis* PrkC in later stages of bacterial growth and germination [41-44]. Even though *prkC* is expressed maximally during the logarithmic phase of *in vitro* growth, *prkC* deletion has never been reported to result in any apparent defect in morphology, viability, or growth during this phase in either *B. subtilis* or *B. anthracis* [38-41, 43, 44, 50]. PrkC is, however, recognized as an infection-specific kinase and is critical for *B. anthracis* survival in macrophages [39, 40].

While working on the *B. anthracis* Sterne 34F2 *prkC* mutant strain (BAS Δ*prkC*), we observed that logarithmic cultures of BAS Δ*prkC* allowed to stand at room temperature formed a compact pellet whereas the parental wild type strain (BAS WT) did not (Fig 1A). The absence of PrkC was leading to the formation of a compact pellet. In a study on a *B. anthracis bslO* mutant strain, Anderson et al. had shown that compact pellets were formed when bacteria grew as shorter chains while loose pellets were formed when bacteria exhibited extensive chaining [28]. This suggested that the absence of PrkC might be leading to the shortening of bacterial chains. To validate this, exponentially growing BAS WT and BAS Δ*prkC* were visualized under a phase-contrast microscope. As shown in Fig 1B, disruption of *prkC* resulted in bacteria growing as shorter chains, and this phenotype was reversed in a *prkC*-complemented strain (BAS Δ*prkC*::*prkC*). Further, to determine if *prkC* disruption resulted in a defect in the cell morphology, viz; bulging, shrinking, or changes in cell width or shape, BAS WT, and BAS Δ*prkC* were examined by scanning electron microscope (SEM). However, as seen in Fig 1C, no morphological defect was apparent, apart from the shortening of bacterial chains, indicating that the *prkC* disruption influenced only chain length. In a study on PknB, a membrane-localized PASTA kinase from *Mycobacterium tuberculosis*, depletion, or over-expression of the kinase was shown to have a significant effect on bacterial morphology leading to cell death [51].

**Fig 1.**
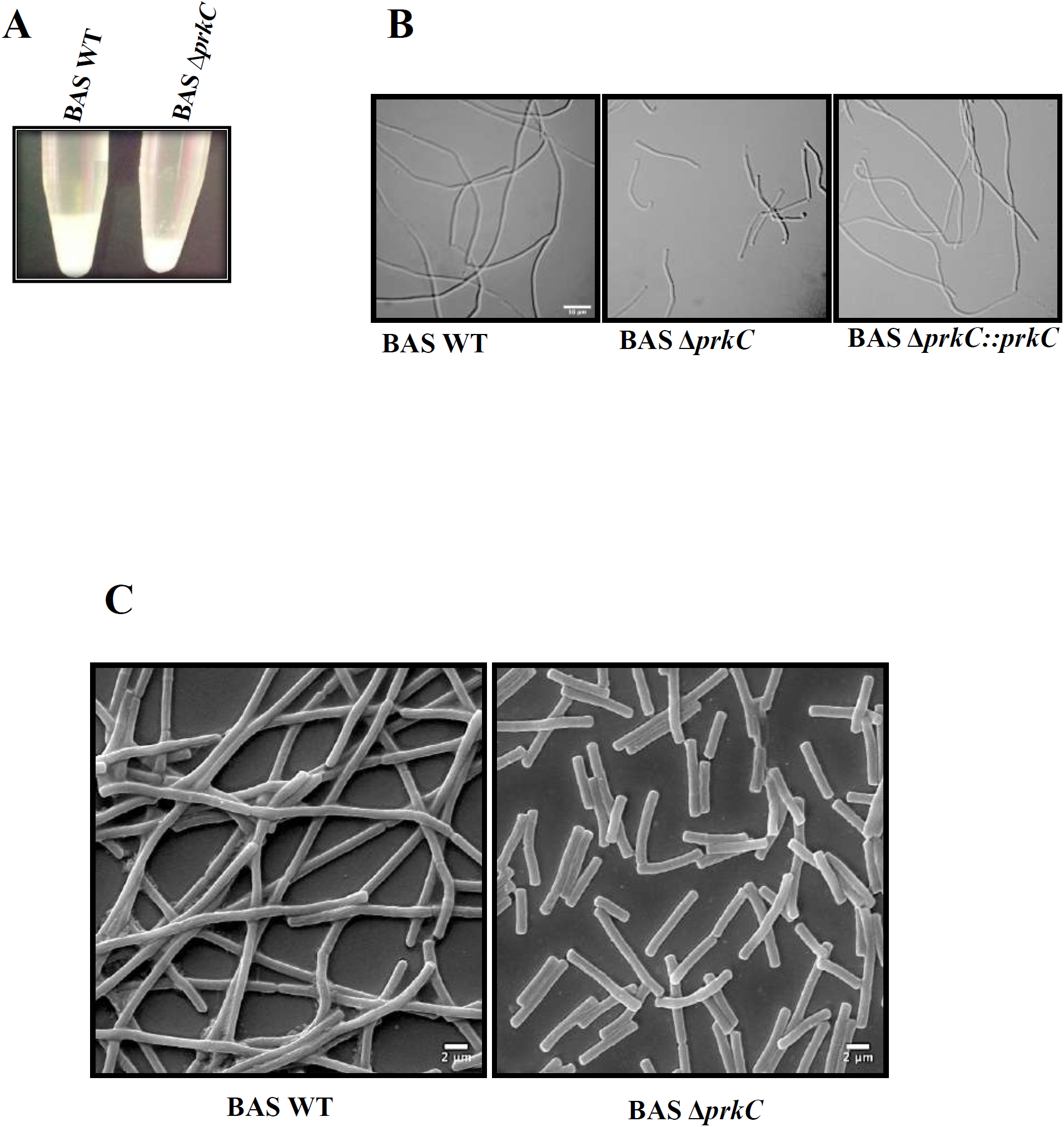
*prkC* disruption results in bacteria with short chain length. (A) Photograph of culture sediments in microcentrifuge tubes after standing incubation (9 hr) at room temperature of BAS WT (left) and BAS Δ*prkC* (right) grown in LB media. (B) Phase contrast images of BAS WT, BAS Δ*prkC* and BAS Δ*prkC*::*prkC* strains in mid-log phase. Cells were grown in LB broth at 37°C and 1 ml sample was taken from cultures in mid-log phase. Cells were pelleted and washed with PBS and visualised under 100x/1.4 oil DIC objective of Zeiss Axio Imager Z2 Upright Microscope. Scale bar represents 10 μm. (C) Scanning electron microscopy of BAS WT and BAS Δ*prkC* strains in mid-log phase. Cells were grown in LB broth at 37°C and harvested in mid-log phase. These were then washed with 0.1 M sodium phosphate buffer and fixed with Karnovasky’s fixative followed by 1% osmium tetroxide. A critical point drying technique was used for drying the samples followed by gold coating of 10 nm using an aluminium stubs coated with agar sputter. Cells were visualized under Zeiss Evo LS15. Scale bar represent 2 μm, magnification-5000X.

### Effect of *prkC* disruption on chaining morphotype during different phases of bacterial growth

If PrkC is the sensor molecule required for maintaining the chaining phenotype, its absence in the BAS Δ*prkC* strain would result in shorter chains throughout the bacterial growth cycle. To examine this and to provide a basis for our experiments, we first monitored the growth of the BAS WT strain through the entire growth cycle (Fig 2A). To determine growth stage-specific changes in chaining phenotype, culture aliquots were taken out at indicated time points and observed under a phase-contrast microscope. As seen in Figs 2C and 3, BAS WT exhibited extensive chaining until ∼ OD (*A600*_*nm*_) 3.0, at 4 hr, after which a sudden shortening of bacterial chains was observed. The average chain length at 4hr was measured as 115.90 (±46.271 S.D., n = 50) µm while at 5 hr it shortened to 63.53 (±20.1150 S.D., n = 50) µm (Fig 3). Interestingly, this time point correlated with the end of the exponential phase and the start of the deceleration phase (Fig 2A), a stage where bacterial replication rate starts decreasing owing to nutrient deprivation and accumulation of metabolic by-products [52]. Next, to determine the effect of *prkC* disruption on chaining phenotype, similar growth curve analysis, and chain length determinations/measurements were carried out as described above (Figs 2B, 2C, and 3). As shown in Figs 2C and 3, BAS Δ*prkC* grew as shorter chains throughout the growth cycle. Of note, we did observe some chaining in the BAS Δ*prkC* cultures during the lag phase (t = 2h, Fig 2C), which could be due to an insufficient number of the de-chaining molecule(s) synthesized at this stage. In the presence of PrkC, synthesis of these molecule(s) is probably downregulated to allow bacteria to grow as chains. These results indicate that PrkC senses growth permissive signal(s) and regulates the levels of molecules associated with de-chaining to maintain the long-chain phenotype, a morphology found during nutrient abundance [3, 28].

**Fig 2.**
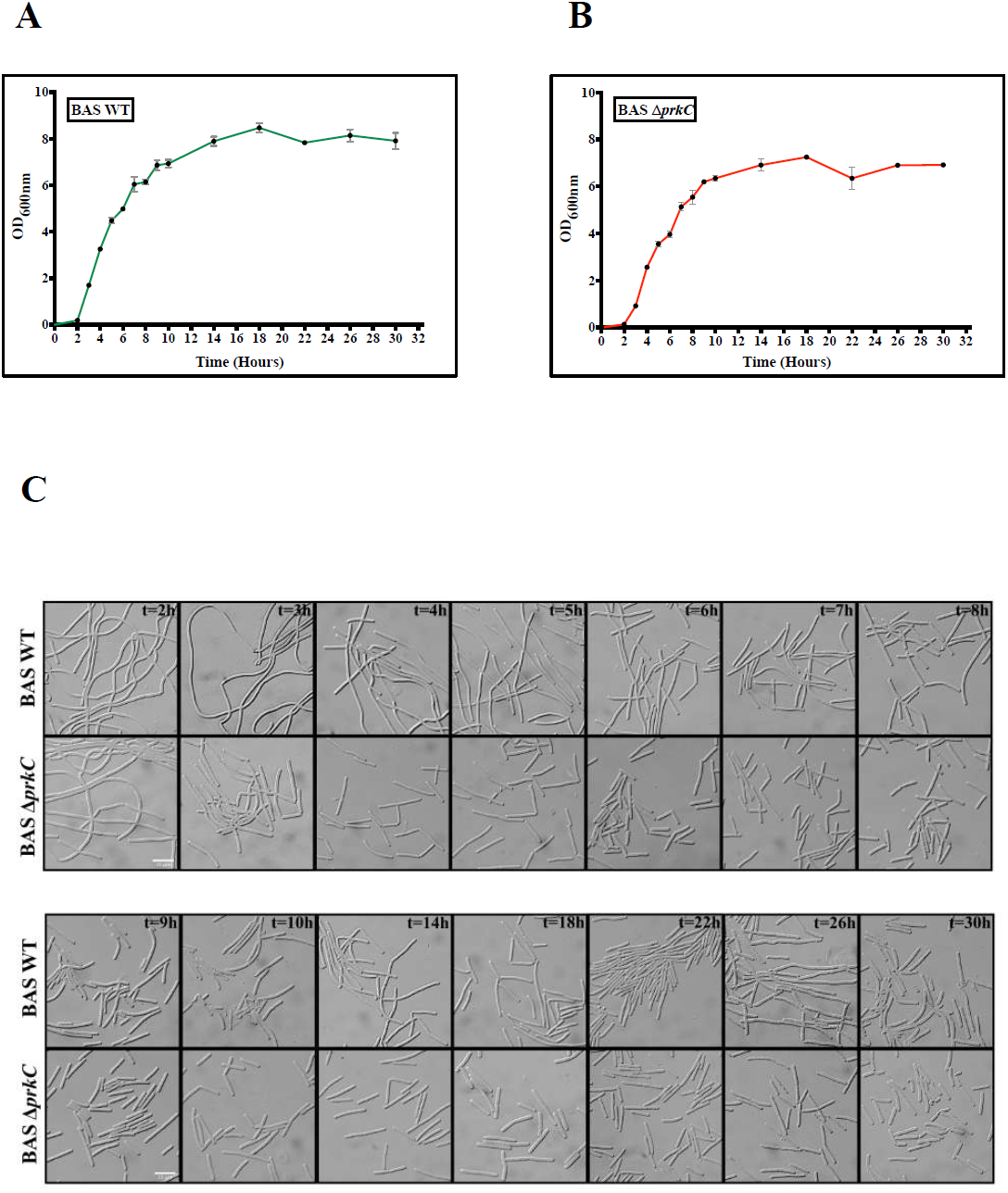
Effect of *prkC* disruption on chaining morphotype during different phases of bacterial growth. (A) Growth kinetics of BAS WT. BAS WT strain was grown in LB broth at 37°C. Absorbance [OD (*A600*_*nm*_)] was recorded at the indicated time points. Error bars denote standard deviation, n = 3. (B) Growth kinetics of BAS Δ*prkC*. BAS Δ*prkC* strain was grown in LB broth at 37°C. Absorbance [OD (*A600*_*nm*_)] was recorded at the indicated time points. Error bars denote standard deviation, n = 3. (C) Phase contrast images of BAS WT and BAS Δ*prkC* strains at different phases of bacterial growth cycle. Cells were grown at 37°C in LB broth and 1 ml sample was harvested at time points indicated in Fig. 2A and Fig. 2B. Cells were pelleted and washed with PBS and visualised under 100x/1.4 oil DIC objective of Zeiss Axio Imager Z2 Upright Microscope. Scale bar represents 10 μm.

**Fig 3.**
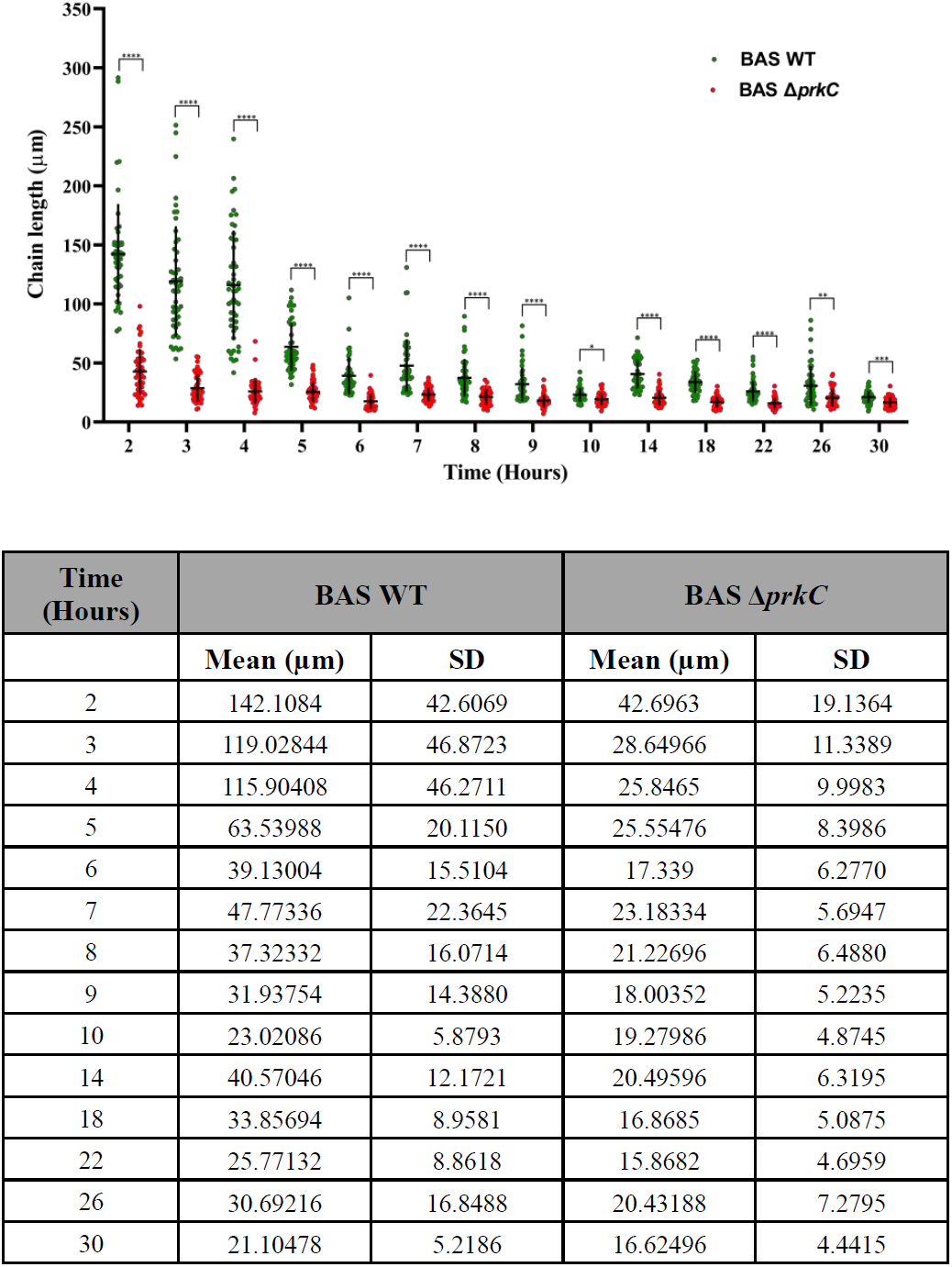
Quantitative analysis of chain length variation in *prkC* disruption strain. Scatter dot plot denoting BAS WT and BAS Δ*prkC* strain chain length measurement throughout bacterial growth. Phase contrast images of BAS WT and BAS Δ*prkC* strains were used for measurement of the bacterial chain length using ImageJ software. Some data points in chain length quantitation (BAS WT – 2 hr, 3 hr and 4 hr) represent the maximum observable chain length obtained in the phase contrast images. Vertical and horizontal black line in the data set denotes SD and mean for both the strains at indicated time points, n = 50. These values are indicated in a separate table below the graph. Statistical significance of chain length distribution in BAS WT and BAS Δ*prkC* strains was analyzed using two-way ANOVA and denoted in the graph in the form of asterisk - *. P-values reported - *, p<0.05; **, p<0.01; ***, p<0.001 and ****, p<0.0001 were considered significant results. Detailed summary statistics table is provided in supporting information - S4 Table.

### PrkC regulates the expression of Sap, EA1 and BslO

In *B. anthracis*, Sap, and then EA1 sequentially form monomeric paracrystalline bi-dimensional surface S-layers during exponential and stationary growth-phase, respectively [27, 53]. The saturating presence of Sap and EA1 on the cell wall confines the S-layer associated protein BslO (with a similar SLH domain) to the septal region [27]. Disruption of *sap* has been shown to cause chain length elongation mainly because in the absence of Sap, BslO is no longer restricted to the septal region and is hence incapable of carrying out murein hydrolysis effectively [27]. To understand whether PrkC maintains chaining phenotype through modulating the levels of Sap, BslO, and EA1, their expression levels were determined at the indicated time points (Figs 2A and 2B) in the BAS WT and BAS Δ*prkC* strains. As shown in Fig 4A, *prkC* disruption resulted in the upregulation of Sap at most of the time points for which the samples were collected. In BAS WT, a sharp increase in expression was observed toward the end of the exponential growth phase {∼ OD (*A600*_*nm*_) 3, Time – 4 hr}, Figs 2A and 4A. Interestingly, the initial time points when the Sap expression was low were also the time points where long chains were observed (Time – 2 hr and 3 hr), Figs 2C, 3 and 4A. However, in BAS Δ*prkC* strain, Sap levels were found to be higher than the BAS WT strain even at the initial time points (Time – 2 hr and 3 hr), Fig 4A. This probably formed the reason for the de-chaining observed in the BAS Δ*prkC* strain from the beginning of the growth cycle. These results are in agreement with the previous report, where the absence of Sap was shown to result in a long chain phenotype [27].

**Fig 4.**
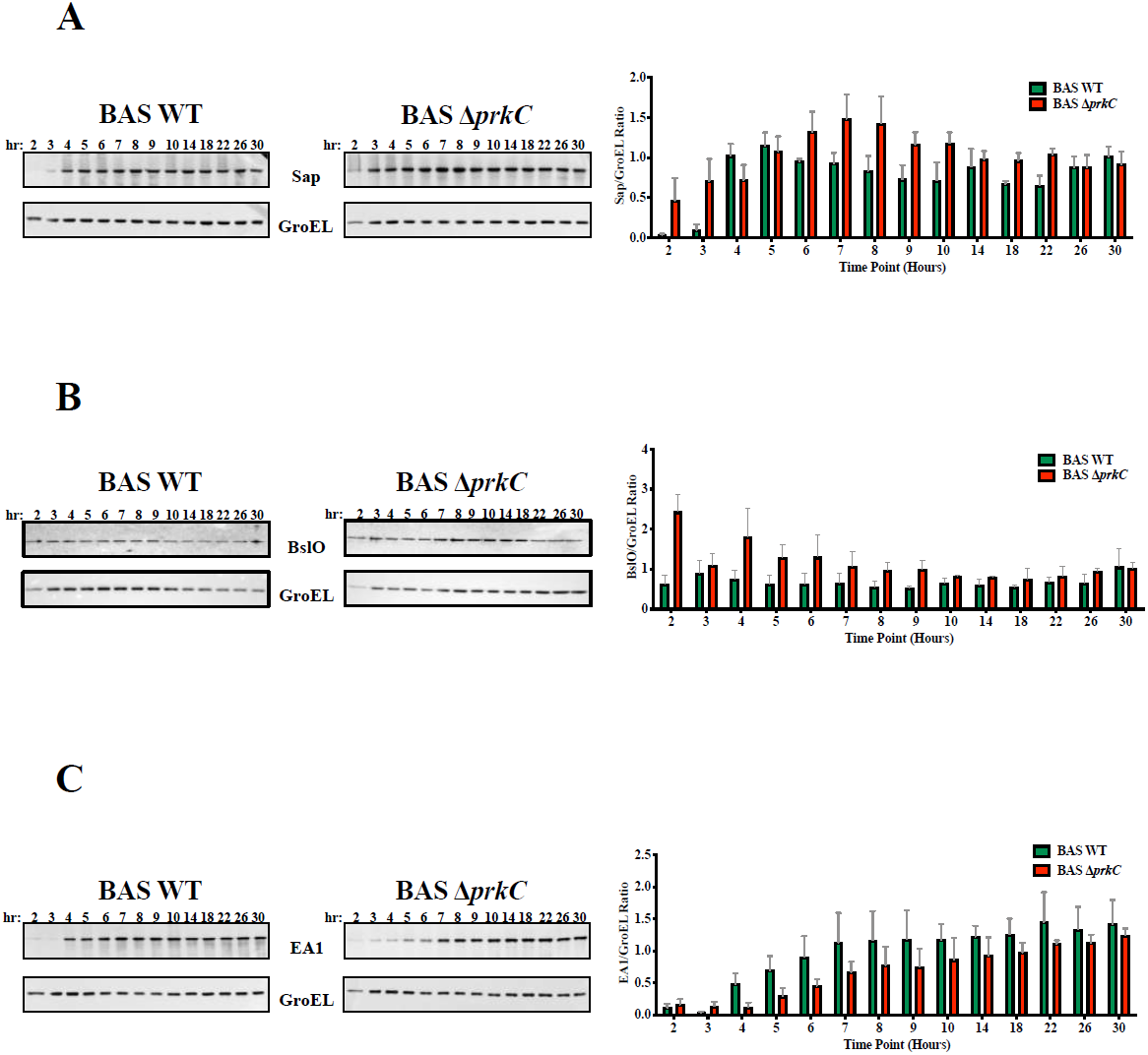
PrkC regulates the expression of Sap, BslO and EA1. (A) Differential expression of Sap protein in BAS WT and BAS Δ*prkC* strains. Equal amount of protein at different time points (2 hr, 3 hr, 4 hr, 5 hr, 6 hr, 7 hr, 8 hr, 9 hr, 10 hr, 14 hr, 18 hr, 22 hr, 26 hr and 30 hr) was loaded onto SDS-PAGE, transferred onto nitrocellulose membrane and probed by Sap antibody (1:50,000) raised in rabbit. The same blot was then stripped and probed by GroEL antibody (1:50,000) raised in mice. Densitometry analysis was done using ImageLab software and the ratio of Sap w.r.t. GroEL was used to plot a graph for showing differential expression of Sap protein in the BAS WT and BAS Δ*prkC* strain throughout bacterial growth. Error bars denote standard deviation, n = 3. Representative images are from one of the three independent experiments. (B) Differential expression of BslO protein in BAS WT and BAS Δ*prkC* strains. Equal amount of protein at different time points (2 hr, 3 hr, 4 hr, 5 hr, 6 hr, 7 hr, 8 hr, 9 hr, 10 hr, 14 hr, 18 hr, 22 hr, 26 hr and 30 hr) was loaded onto SDS-PAGE, transferred onto nitrocellulose membrane and probed by BslO antibody (1:10,000) raised in mice. The same blot was then stripped and probed by GroEL antibody (1:50,000) raised in mice. Densitometry analysis was done using the ImageLab software and the ratio of BslO w.r.t. GroEL was used to plot a graph for showing differential expression of BslO protein in the BAS WT and BAS Δ*prkC* strain throughout bacterial growth. Error bars denotes standard deviation, n = 3. Representative images are from one of the three independent experiments. (C) Differential expression of EA1 protein in BAS WT and BAS Δ*prkC* strains. Equal amount of protein at different time points (2 hr, 3 hr, 4 hr, 5 hr, 6 hr, 7 hr, 8 hr, 9 hr, 10 hr, 14 hr, 18 hr, 22 hr, 26 hr and 30 hr) was loaded onto SDS-PAGE, transferred onto nitrocellulose membrane and probed by EA1 antibody (1:50,000) raised in rabbit. The same blot was then stripped and probed by GroEL antibody (1:50,000) raised in mice. Densitometry analysis was done using ImageLab software and the ratio of EA1 w.r.t. GroEL was used to plot a graph for showing differential expression of EA1 protein in the BAS WT and BAS Δ*prkC* strain throughout bacterial growth. Error bars denotes standard deviation, n = 3. Representative images are from one of the three independent experiments.

Next, we wanted to determine the levels of BslO in the BAS WT and BAS Δ*prkC* strains. Experiments were conducted in a similar manner, as described above. Interestingly, we observed a stable expression of BslO throughout the growth cycle in BAS WT (Fig 4B). As per our understanding, this is the first report where the levels of BslO have been determined at various stages of bacterial growth. Previous studies have conclusively established the role of BslO in de-chaining and have identified it as the primary murein hydrolase driving the de-chaining process [27, 28]. As observed for Sap, *prkC* disruption resulted in the upregulation of BslO at most of the time points for which the samples were collected (Fig 4B). The results obtained for Sap and BslO indicated that an increase in the levels of Sap and BslO in the absence of PrkC might create a condition that is most suitable for de-chaining. Increased Sap would restrict BslO to the septal region, which would carry out de-chaining, and an increase in the levels of BslO would further add to this effect.

Further, we determined the levels of EA1, another structural S-layer protein, that shows its presence as the culture approaches the stationary phase [27, 53]. As seen in Fig 4C, EA1 levels in BAS Δ*prkC* were downregulated at most of the time points for which the samples were collected. Since Sap acts as a transcriptional repressor of *eag* [27, 53], this decrease can be due to the increased levels of Sap in the BAS Δ*prkC* strain (Fig 4A).

Altogether, these results indicate that during bacterial growth, PrkC maintains an optimum level of BslO, Sap, and EA1 to maintain the chaining phenotype.

### *prkC* disruption results in decreased cell wall width and cell septa thickness and increased multi-septa formation

Through the course of these experiments, we observed that BAS Δ*prkC* growth curve was not superimposable with BAS WT (Figs 2A and 2B). This result was in contradiction with the earlier reports wherein *B. anthracis prkC* disruption was shown have no effect on the bacterial growth *in vitro* [39, 40, 48]. To validate our observation, we carried out a comparative growth curve analysis with BAS WT and BAS Δ*prkC* until extended stationary phase. Interestingly, as shown in Fig 5A, BAS Δ*prkC* strain showed an attenuated replication rate throughout the bacterial growth cycle. To identify the reason(s) for the observed defect, we carried out microscopic analysis at the ultrastructural level, and both BAS WT and BAS Δ*prkC* were subjected to transmission electron microscopy. Interestingly, at the ultrastructural level, an apparent decrease in the cell wall width and septal thickness was observed in the *prkC* disruption strain (mid-log phase) (Figs 5B and 5C). Additionally, we also observed an increase in multi-septa formation in the *prkC* disruption strain during later stages of bacterial growth through both confocal and transmission electron microscopy (Figs 6A, 6B and 6C). PASTA domain containing kinases from other bacterial species have been shown to play a role in the regulation of cell division machinery and cell wall homeostasis [54]. We surmise that similar signaling mechanisms may be operational in *B. anthracis* as well.

**Fig 5.**
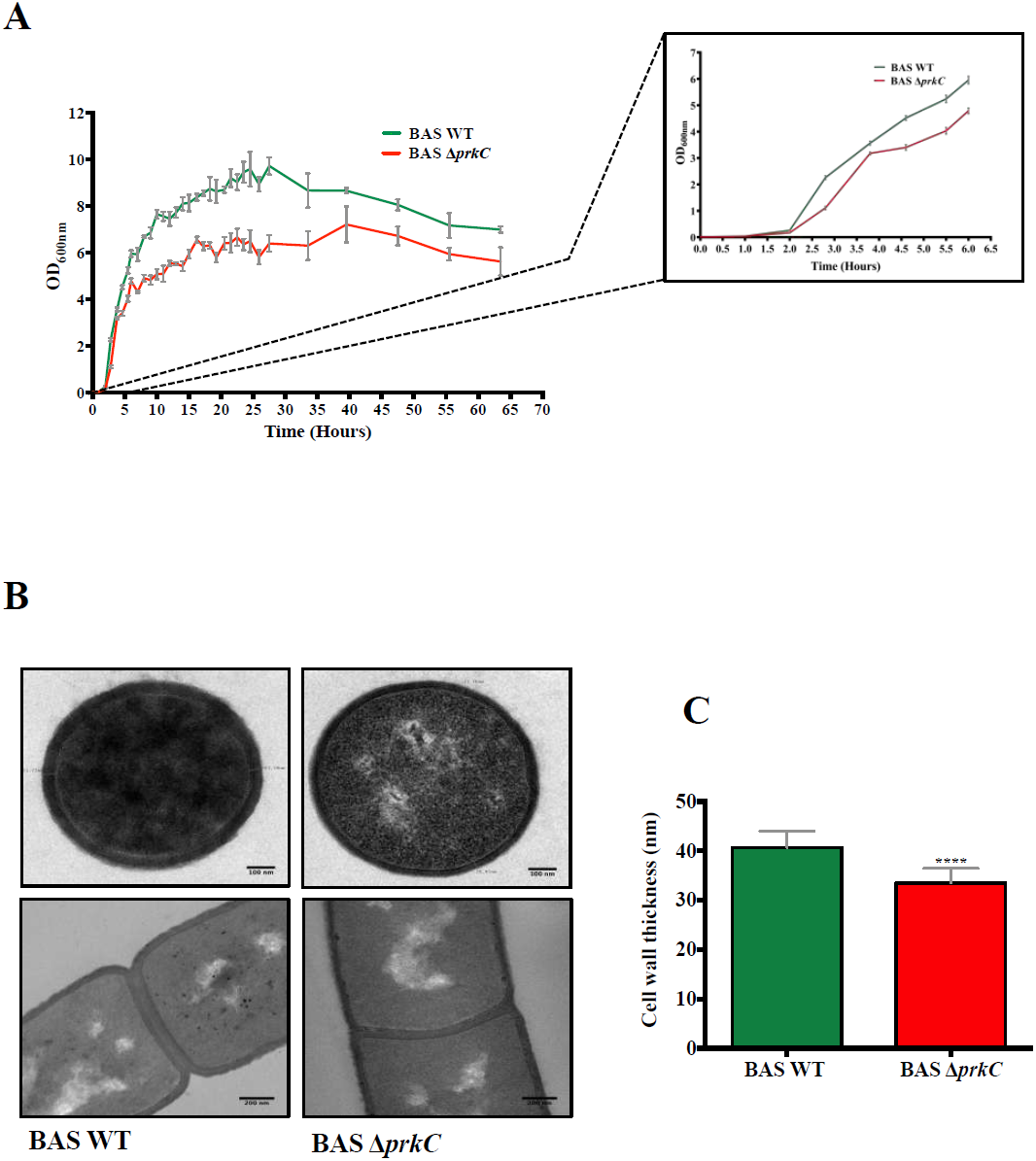
*prkC* disruption results in decreased cell wall width and septa thickness. (A) Growth kinetics of BAS WT and BAS Δ*prkC* strains. Bacterial strains were grown in LB broth at 37°C till extended period of 65 hr. Absorbance [OD (*A600*_*nm*_)] was recorded at the indicated time points. Error bars denote standard deviation, n = 3. Inset shows expanded growth profile of BAS WT and BAS Δ*prkC* strains up till 6 hr. (B) Transmission electron micrographs representing the ultrastructural details of difference in cell wall thickness and septum thickness of BAS WT and BAS Δ*prkC* strains. Cells were harvested at mid-log phase and primary fixation was done using Karnovasky’s fixative. Secondary fixation was done using 1% osmium tetroxide and the samples were embedded into araldite resin mixture (TAAB). Scale bar represents 100 nm. (C) Bar graph representing the difference in cell wall thickness of BAS WT and BAS Δ*prkC* strains. Transverse sections of around 100 mid-log phase cells of each strain was used to calculate the cell wall thickness and plotted. Statistical significance of the data set was analysed using two-tailed Student’s *t* test and denoted in the graph in the form of asterisk - *. P-values reported - *, p<0.05; **, p<0.01; ***, p<0.001 and ****, p<0.0001 were considered significant results. Detailed summary statistics table for Fig 5C is provided in supporting information - S5 Table.

**Fig 6.**
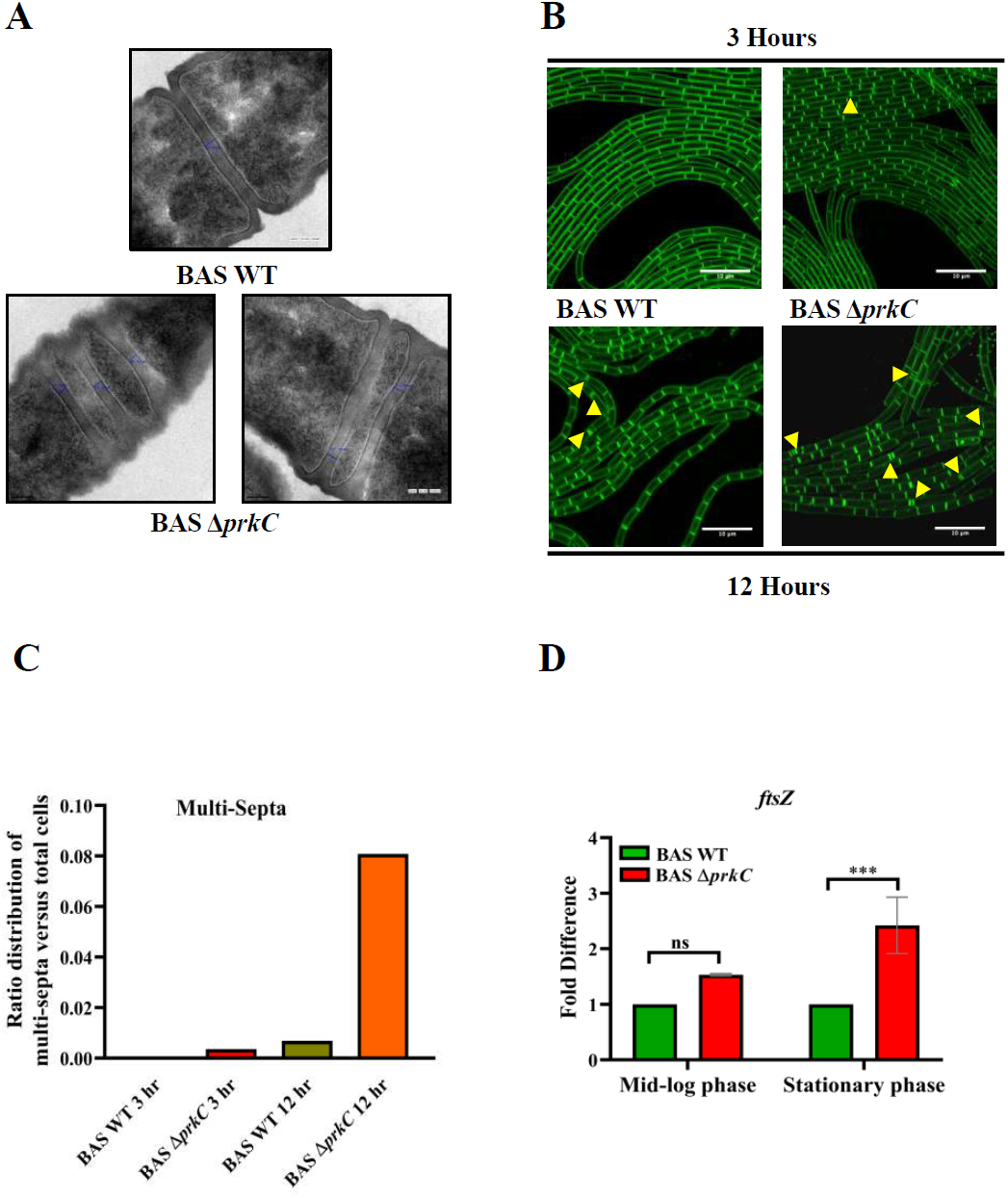
*prkC* disruption results in increased multi-septa formation. (A) Representative transmission electron microscopy images of BAS WT and BAS Δ*prkC* stationary phase cells showing multi-septa formation. Cells were harvested at stationary phase and primary fixation was done using Karnovasky’s fixative. Secondary fixation was done using 1% osmium tetroxide and the samples were embedded into araldite resin mixture (TAAB). Scale bar represents 100 nm. (B) Staining of live bacterial cells with FM4-64 membrane stain. BAS WT and BAS Δ*prkC* strains grown up to exponential phase were diluted to an initial [OD (*A600*_*nm*_)] = 0.035 and 1μL was spread on agarose pad. The pads were incubated at 37°C. Cell membrane was stained with FM4-64 (Final concentration of 1 μg/ml) and images were captured by Leica SP8 confocal microscope at 3 hr (above panel) and 12 hr (below panel). Arrows indicate the presence of multi-septa in the images. Scale bar represents 10μm. (C) Graph indicating ratio distribution of multi-septa formation with respect to the total number of cells in BAS WT and BAS Δ*prkC* strains at 3 hr and 12 hr. Around 1500 cells were considered for calculation of each bar in the graph. (D) Comparative gene expression analysis of *ftsZ* gene in BAS Δ*prkC* strain as compared to BAS WT strain during mid-log and stationary phase. The data was normalized to the expression of *rpoB* from each sample. Error bar represents an average of three biological and three technical replicates. Statistical significance of *ftsZ* gene expression in BAS WT and BAS Δ*prkC* strains at both the time points was analyzed using two-way ANOVA and denoted in the graph in the form of asterisk - *. P-values reported - *, p<0.05; **, p<0.01; ***, p<0.001 and ****, p<0.0001 were considered significant results. Detailed summary statistics table for Fig 6D is provided in supporting information - S6 Table.

FtsZ is a cytoskeletal protein of cell division machinery that localizes at mid-cell and forms the initial Z ring. It also serves as the scaffold for further assembly of cell-division machinery [55]. STPKs from other bacterial systems have been shown to phosphorylate and regulate the activity of FtsZ [56, 57]. Our initial results suggest that *ftsZ* is constantly upregulated in the *prkC* disruption strain (Fig 6D). This probably formed the reason for an increase in the multi-septa formation observed in the *prkC* disruption strain, possibly due to mis-localization of FtsZ. Further experiments are underway to delineate the PrkC-mediated signaling cascades, disruption of which results in the observed defects in cell division, cell wall homeostasis, and multi-septa formation.

## Discussion

Chaining phenotype acts as a virulence factor in several bacterial pathogens [1, 4, 58-63]. In *Legionella pneumophila*, chaining morphology helps the pathogen evade phagosomal killing by interfering with phagosomal morphogenesis [60]. In *Streptococcus pneumoniae*, bacteria growing as long chains display increased attachment and adherence to epithelial cell surfaces, possibly via multivalent binding sites [59]. *Bacillus cereus*, a close relative of *B. anthracis* and a cause of food-borne and opportunistic infections in humans, also displays a chaining phenotype and has been shown to attach to the invertebrate gut through long filaments [62, 63]. In *B. anthracis*, in a mice model, chain-length-dependent physical sequestration of an encapsulated nontoxinogenic strain in lung capillaries is thought to result in hypoxia and associated lung tissue injury, leading to host death [4, 5].

In this study on *B. anthracis* Sterne strain, we identify a transmembrane serine/threonine protein kinase, PrkC, with an extracellular sensory PASTA domain, as a determinant of chaining phenotype. Interestingly, PrkC homologs are found in all the above-mentioned chain-forming pathogens (S1 Fig). Though PASTA kinases do not necessarily carry out similar signaling processes across various bacterial species [37, 64], it would be worthwhile to explore whether PASTA kinases of these pathogens also form the primary messenger molecule for maintaining chaining phenotype as reported for *B. anthracis* in this study. Notably, PASTA motifs are unique to bacteria, and their absence in eukaryotes makes them an attractive drug target [64].

*B. anthracis* PrkC is a key messenger molecule that plays a central role in various cellular processes including infection in macrophages, biofilm formation and germination [39, 40, 42, 48]. We report for the first time that *prkC* disruption results in the attenuation of growth rate during *in vitro* culturing (Fig 5A). Interestingly, none of the previous studies have reported an attenuation of growth on the disruption of *prkC* during *in vitro* growth, as observed in this study. We believe that this discrepancy could be due to the difference in the subtype of the Sterne strain used [*B. anthracis* Sterne strain 7702 [39, 40] vs. B. *anthracis* Sterne strain 34F2 {used in this study}] or differential growth conditions.

In this study, we show that in the *prkC* disruption strain, S-layer protein, Sap, and septal *N*-acetylglucosaminidase, BslO, are upregulated. On the contrary, stationary phase S-layer protein, Ea1, shows downregulation (Fig 4). S-layers are found in many bacterial species where they form a cell cover and play important roles such as - 1) act like exoskeleton/mechanical barrier, 2) function like a scaffold for surface molecules, 3) mediate adhesion, leading to auto-aggregation and coaggregation, 4) work as a molecular sieve and, 5) act as an immunomodulatory factor [65, 66]. In *B. anthracis*, S-layer is made up of two primary structural proteins, Sap and EA1, and several S-layer-associated proteins called BSLs, that carry out diverse roles [67]. Previous studies have shown that Sap levels rise until the onset of the stationary phase, and the EA1 amount is minimal during the logarithmic phase [53]. As Sap levels go down, EA1 is upregulated and replaces Sap as the primary constituent of S-layer in the stationary phase. Both Sap and EA1 act as the transcriptional repressors of the *eag* gene [53]. Our results also show a gradual decline in the Sap levels from late log phase onwards {∼ OD (*A600*_*nm*_) 5.0-6.0, Time – 6-8} (Figs 4A and 2A). EA1 levels, as reported earlier, were minimal during lag phase and early exponential phase but increased gradually till the last point of measurement (Fig 4C). Interestingly, BslO levels remained constant throughout the growth cycle in BAS WT (Fig 4B), which implies that the stage/growth phase-dependent de-chaining in wild-type strain is primarily dependent on its localization, which in turn is controlled by the levels of Sap on the cell surface. Upregulation of both Sap and BslO in BAS Δ*prkC* strain would create a condition that would favor de-chaining from the initial stages of bacterial growth. Altogether, our results suggest that PrkC keeps a check on the levels of Sap, BslO, and Ea1 during optimum growth conditions, thereby maintaining the chaining phenotype.

In conclusion, through this study, we show that PrkC, the transmembrane kinase of *B. anthracis* with a sensor PASTA domain, regulates chaining phenotype. Since the disruption strain of PrkC shows decreased virulence in mice model of pulmonary anthrax [39], it will be relevant to see if the observed effect is due to a difference in the chaining phenotype as shown in this report. If proven so, therapeutic intervention against PrkC could help in controlling bacterial chain size and hence the lung tissue injury and its pathophysiological consequences.

## Materials and methods

### Bacterial strains and growth conditions

*Escherichia coli* DH5α (Invitrogen) and SCS110 (Stratagene) strains were used for cloning and BL21-DE3 (Invitrogen) strain was used for expression of recombinant proteins. The final concentrations of the antibiotics used were: 100 μg/ml ampicillin, 25 μg/ml kanamycin and 150 μg/ml spectinomycin. LB broth (Difco) with appropriate antibiotic was used to grow bacterial cultures at 37°C with proper aeration (1:5 head space) and constant shaking at 200 rpm. *B. anthracis* Sterne strain 34F2 (BAS WT) was obtained from Colorado Serum Company and *prkC* gene knockout strain (BAS Δ*prkC*) was a gift from Jonathan Dworkin, Department of Microbiology, Columbia University, USA. The details of the primers, plasmids and strains used in the study are provided as supporting information (S1 Table – S3 Table).

### Generation of *prkC* complement strain

The *prkC* gene and its promoter gene sequence were amplified using the primers – (P13-P16). These genes were subsequently cloned in the shuttle vector pYS5 [68] and the positive clone obtained was transformed in SCS110 cells prior to electroporation in the BAS Δ*prkC* strain using Bio-Rad Gene Pulser Xcell (2.5 kV, 400Ω, 25 µF using 0.2 cm Bio-Rad Gene Pulser cuvette). The complemented strain thus obtained after the screening was named BAS Δ*prkC::prkC*.

### Cloning, gene expression, and protein purification

Genes for *sap, eag, bslO* and *groEL* were amplified using BAS genomic DNA as template and sequence specific primers (P5 and P6 - *sap*, P7 and P8 - *eag*, P9 and P10 - *groEL*, P11 and P12 - *bslO*). The amplified products thus obtained for *sap, eag* and *groEL* were cloned in pProExHtc vector (Invitrogen), while *bslO* gene was cloned in pET28a vector (Invitrogen). The resulting plasmids encodes His_6_ tagged fusion proteins. Plasmids were transformed into *E. coli* BL21 (DE3) and proteins were purified using affinity chromatography, as described described [69]. Briefly, overnight grown cultures were diluted in LB broth (1:50) with appropriate antibiotic - ampicillin (100 μg/ml) or kanamycin (25 μg/ml) and grown at 37°C, 200 rpm. Cultures were induced with 1 mM IPTG at OD (*A600*_*nm*_) of 0.5 - 0.8 and incubated overnight at 16°C. After this cells were pelleted and re-suspended in sonication buffer [50 mM Tris-HCl (pH-8.5), 5 mM – β-mercaptoethanol, 1 mM Phenylmethylsulfonyl fluoride (PMSF), 1X protease inhibitor cocktail (Roche Applied Science, U.S.A.) and 300 mM NaCl] and sonicated (9 cycles-20% amplitude, 10 sec on and 30 sec off). The recombinant proteins were purified by affinity purification using Ni-nitrilotriacetic acid (NTA) column and the final elution was done using 200 mM imidazole. Protein estimation was done using Pierce BCA Protein Assay kit (Thermo Fisher Scientific).

### Generation of polyclonal antibodies against GroEL and BslO in mice, and Sap and EA1 in rabbit

For generation of polyclonal antibodies 30 μg of purified protein (GroEL and BslO) was used for injecting in three BALB/c mice and 500 μg of purified protein (Sap and EA1) was used for injecting in three rabbits for each protein. Antigens were emulsified in complete Freund’s adjuvant (Sigma-Aldrich) in 1:1 ratio before subcutaneous injection in the animals. Production of antibody was stimulated at an interval of 21 days followed by two booster injections of 15 μg protein emulsified in incomplete Freund’s adjuvant for mice and three booster injections of 250 μg protein emulsified in incomplete Freund’s adjuvant for rabbits. Animals were bled to collect serum 14 days after the final injection and the antibody titer was calculated by ELISA and used accordingly for further experiments.

### Growth Kinetics

BAS WT and BAS Δ*prkC* strains were grown overnight in LB broth at 37°C, 200 rpm. These overnight grown cultures were taken as inoculum for growth kinetics experiments. The secondary cultures were initiated at starting OD (*A600*_*nm*_) of 0.001 in triplicates in LB broth at 37°C, 200 rpm (New Brunswick Innova 42 Incubator Shaker). Following this the OD (*A600*_*nm*_) was monitored until around 64 hr at indicated intervals (Fig 5A).

For lysate preparation and microscopy, the secondary cultures for both the strains were initiated similarly in triplicates from overnight grown cultures at starting OD (*A600*_*nm*_) of 0.001. Culture samples were collected at different time points (2 hr, 3 hr, 4 hr, 5 hr, 6 hr, 7 hr, 8 hr, 9 hr, 10 hr, 14 hr, 18 hr, 22 hr, 26 hr, and 30 hr) for analysis and the corresponding OD (*A600*_*nm*_) was plotted (Fig 2A and 2B).

### Phase Contrast Microscopy

For phase-contrast microscopy 1 ml of culture samples from growing cells of BAS WT and BAS Δ*prkC* were collected at different time points as mentioned in above section. The cells were pelleted and washed thrice with phosphate buffer (pH 7.4) and resuspended in 100 μL buffer. The cells were then observed under a 100 x/1.4 oil DIC objective of Zeiss Axio Imager Z2 Upright Microscope. Images were captured using Axiocam 506 color camera equipped to the microscope and processed in ZEN 2 Pro software.

### Quantitative Immunoblot Analysis

For immunoblot analysis, 5-10 ml bacillus culture was collected at different time points (2 hr, 3 hr, 4 hr, 5 hr, 6 hr, 7 hr, 8 hr, 9 hr, 10 hr, 14 hr, 18 hr, 22 hr, 26 hr, and 30 hr) from BAS WT and BAS Δ*prkC* cultures. The cell pellet obtained was washed with PBS buffer and re-suspended in 1 ml lysis buffer [200 mM Tris-HCl (pH 7.5), 1 mM EDTA, 150 mM NaCl, 1 mM PMSF, 5% glycerol, and 1X protease inhibitor cocktail (Roche Applied Science)]. After this, sonication of the bacterial pellets resuspended in the buffer was done (9 cycles-20% amplitude, 10 sec on and 30 sec off). Protein estimation was done using Pierce BCA Protein Assay kit (Thermo Fisher Scientific). 5 μg protein lysate sample of each time point was prepared using SDS sample buffer containing 250 mM Tris-HCl (pH 6.8), 30% (v/v) glycerol, 10% SDS, 10 mM DTT and 0.05% (w/v) Bromophenol Blue. Samples were heated for 5 min at 95°C prior to loading on 12% polyacrylamide gels followed by transfer onto NC membrane (Millipore). 3% BSA in phosphate buffer saline (PBS) with 0.05% Tween 20 (PBST) was used for blocking the membranes overnight at 4°C. This was followed by washing with PBST (3 washes of 5 mins each). Membranes were then probed with antibodies specific to Sap protein (1:50,000) or BslO protein (1:10,000) or EA1 protein (1:50,000) for 1 hr followed by washes with PBST (5 washes of 5 min each). After this anti-rabbit IgG secondary antibody (for Sap and EA1) or anti-mouse IgG secondary antibody (for BslO) conjugated with horseradish peroxide (1:10,000 - Cell Signaling Technology) was used and the blots were incubated for another 60 min, followed by 3 PBST washings of 10 min each. Finally, SuperSignal West Pico PLUS Chemiluminescent substrate (Thermo Fisher Scientific) was used to detect the signal and it was visualised and quantified with the luminescent image analyser (Amersham Imager 600 or ImageLab6.0.1). The blots were then stripped using stripping buffer and probed similarly using anti-GroEL (1:50,000) and anti-mice IgG secondary antibody conjugated with horseradish peroxide (1:10,000 - Cell Signaling Technology) for normalising the loading pattern. GroEL was used as a loading control as PrkC has been shown to modify GroEL without affecting its total expression level [48].

### RNA extraction and Quantitative Real Time PCR

BAS WT and BAS Δ*prkC* strains were grown to mid-log and stationary phase in triplicates for RNA extraction following hot lysis method as described previously [70-72] with a few modifications. Cells were harvested at 6,000 x g for 15 min and the cell pellet was washed once with PBS and resuspended in 500 μL TRIzol^®^ (Invitrogen) and frozen at −80°C until ready for further processing. The frozen samples were thawed in ice and RNA was extracted following the hot lysis method. Briefly the samples were mixed with 400 μL of buffer (50 mM Tris (pH 8.0), 1% SDS and 1mM EDTA) and 400 μL of zirconia beads treated with DEPC water. This suspension was incubated at 65 °C for 15 min with rigorous intermittent vortexing after every 5 min. Suspension was cooled in ice and mixed well with 100 μLof chloroform/ml of TRIzol. Separation of the aqueous phase containing RNA was done by centrifugation at 9500 × g for 15 min at 4 °C. RNA was precipitated from the aqueous phase by adding LiCl_2_ (0.5M) and 3X ice-cold isopropanol followed by 2 hr incubation at −80°C. RNA pellet thus obtained by centrifugation at 16,000 × g for 20 min (4°C) was washed using 70% ethanol (Merck) and resuspended in nuclease-free water after air drying. RNA sample was then treated with DNase (Ambion) to remove any residual DNA contamination (according to the manufacturer’s protocol). RNeasy mini kit (Qiagen) was used to obtain pure RNA using the manufacturer’s protocol. cDNA was prepared using 1 μg of RNA using first-strand cDNA synthesis kit (Thermo Fisher) according to the protocol provided by the manufacturer. To analyse the expression of *ftsZ* gene, 2 μL of cDNA (diluted 10 times) was used for each time point (mid-log and stationary) along with gene-specific primer and SYBR Green master mix (Roche) in a 10 μL reaction according to the manufacturer’s protocol. Reactions were run in triplicates along with no template control in a LightCycler^®^ 480 Instrument II (Roche). The *rpoB* gene encoding for DNA-directed RNA polymerase subunit beta was used as housekeeping control [73]. All the primers used were sequence-specific with a PCR product of 120 bp size.

### Scanning Electron Microscopy

BAS WT and BAS Δ*prkC* strains were grown in LB broth at 37°C and harvested at mid-log phase and processed [74]. Briefly the bacterial culture was harvested at 12,000 x g at 4°C, and the pellet thus obtained was washed thrice using 0.1 M sodium phosphate buffer (pH-7.4). Karnovsky’s fixative (2.5% glutaraldehyde (TAAB) + 2% paraformaldehyde (Sigma) in 0.1 M sodium phosphate buffer pH 7.4 was used to fix the bacterial samples overnight at 4°C. Fixed cells were again washed using sodium phosphate buffer and this step was repeated thrice to remove any residual fixative from the pellet. After this the pellets were again fixed using 1% osmium tetroxide for 20 min at 4°C. Sequential dehydration was then done for 30 min each using a range of ethanol (Merck) (30%, 50%, 70%, 80%(X 2), 90%, 100%(X 3) at 4°C. A critical point drying technique was used for drying the samples followed by gold coating of 10 nm using an aluminium stubs coated with agar sputter. Images were captured using Zeiss Scanning Electron Microscope EVO LS15 at 20 KV. Comprehensive imaging, processing and analysis were performed with Smart SEM software [75].

### Transmission Electron Microscopy

BAS WT and BAS Δ*prkC* strains were grown in LB Broth at 37°C and harvested at mid-log and stationary phase and processed. Briefly the bacterial culture was harvested at 12,000 x g at 4°C, and the pellet thus obtained was washed thrice using 0.1 M sodium phosphate buffer (pH 7.4). Karnovsky’s fixative containing 2.5% glutaraldehyde (TAAB) and 2% paraformaldehyde (Sigma) was made in 0.1 M sodium phosphate buffer pH 7.4 and was used for primary fixation of the cells overnight at 4°C. Fixed cells were again washed using sodium phosphate buffer and this step was repeated thrice to remove any residual fixative from the pellet. After this the pellets were again fixed using 1% osmium tetroxide for 20 min at 4°C. Sequential dehydration was then done for 30 min each using a range of acetone (Merck) (30%, 50%, 70%, 80% (X 2), 90%, 100% (X 3) at 4°C. For the clearing process and removal of dehydrating agent, absolute xylene (Merck) was used and the samples were subjected for bullet preparation using araldite resin mixture (TAAB). Following this infiltration was done by raising the concentration of the embedding medium and lowering the concentration of clearing agents gradually. The final bullets were prepared by curing at 55°C for 24 hr and for 48 hr at 65°C. Sectioning were obtained using Leica UC6 ultra-cut to make the grids which were then observed in FEI Tecnai G^2^ Spirit at 200 KV [75].

### Confocal microscopy to analyse multi septa formation

FM4-64 labelling was used to visualise multi-septa formation in BAS WT and BAS Δ*prkC* strains using fluorescence. LB agarose pads were prepared using AB gene frame (Fischer Scientific; 17*54 mm) on frosted glass slides (Corning Micro slide Frosted; 75*25 mm). To prepare agarose pad 3% low melting agarose (Sigma) were poured on these slides and left for solidification until further use. 1 μL of exponentially growing cultures of BAS WT and BAS Δ*prkC* strains diluted to an OD (*A600*_*nm*_) = 0.035 were spread evenly on the agarose pads along with 1ug/ml FM4-64 dye for staining the cell membrane. Images were captured using Leica TCS SP8 confocal laser scanning microscope at 3 hr and 12 hr using 63x oil immersion objective [76].

### Phylogenetic Analysis

The amino acid sequences of PrkC protein in different pathogens: *Bacillus anthracis* Sterne, *Bacillus cereus, Legionella pneumophilia and Streptococcus pneumoniae* were procured from NCBI. These sequences were aligned using T-coffee tool [77]. These protein sequences were then used for generation of phylogenetic tree by Neighbor Joining analysis conducted by MEGA X [78-80]. The representation of branch lengths is in units of evolutionary distances computed by Poisson correction method [81].

### Statistical analysis

GraphPad Software (Prism 6) was used for all the statistical analyses. The statistical tests are indicated in the figure legends and the corresponding two-tailed *t*-test or ANOVA P-values are reported in the graphs wherever required. *, p<0.05; **, p<0.01; ***, p<0.001 and ****, p<0.0001 were considered significant results. Values indicated in the graphs represent mean ± SD, where n = 3 for both the strains at each time point, unless specified otherwise in the figure legend. Error bars are indicative of SD, n = 3. All the experiments were done in biological triplicates to ensure the reproducibility of the obtained data. Real time experiments (Fig 6D) were done in biological and technical triplicates, while growth kinetics (Fig 2A, 2B and 5A) and western blot analysis (Fig 4) was done using biological triplicates. Separate flasks of the same strains are considered biological replicates, while technical replicates refer to multiple readings of the same sample.

## Acknowledgement

We thank Dr. Hemlata Gautam for technical guidance in scanning electron microscopy, CSIR-IGIB, Delhi and Sandip Arya, Anurag Singh, Raj Girish Mishra and other technical staff from Transmission Electron Microscope facility AIIMS, Delhi. We would also like to thank Dr A.K. Goel, Defence Research & Development Establishment, Gwalior for providing Sap and EA1 antibody. We are grateful to Dr. Anurag Agrawal, Director, CSIR-IGIB for support and providing the required facilities.

## Supporting information

**S1 Fig.**
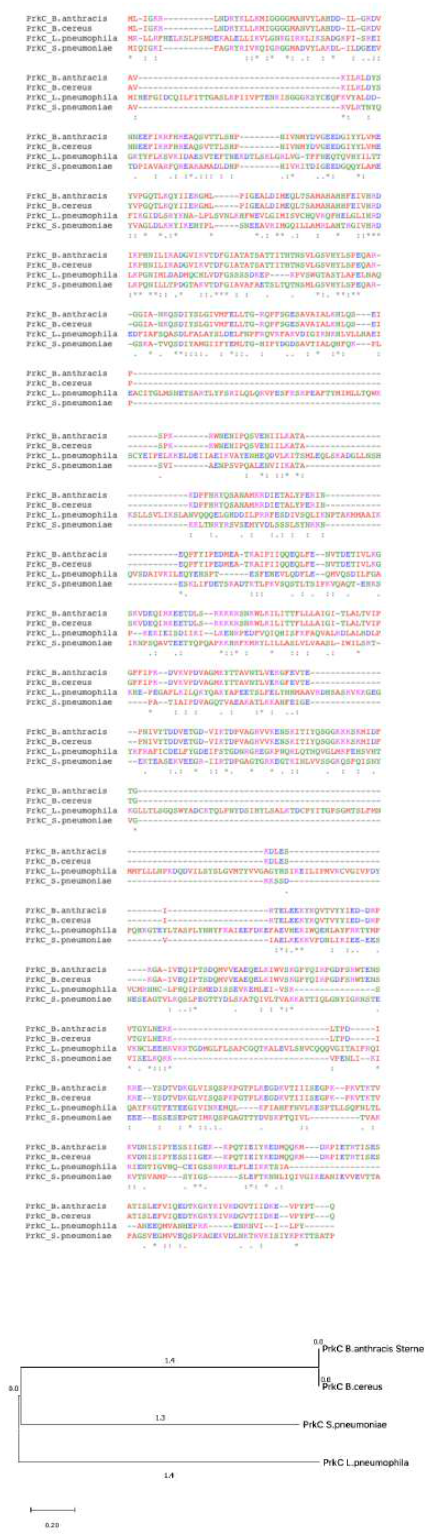
Multiple sequence alignment and phylogenetic analysis of PrkC. **(A)** Multiple sequence alignment of PrkC in different pathogenic bacteria (*Bacillus anthracis* Sterne, *Bacillus cereus, Legionella pneumophilia and Streptococcus pneumoniae)* using T-Coffee program. The symbols used in the fig are indicative of: “**\*”** - perfect alignment, **“:”** - strongly similar residues and **“.”** – weakly similar residues. (B) A phylogenetic tree representing the evolutionary relationship of PrkC in above-mentioned pathogens. It was generated using Neighbor-Joining analysis conducted in MEGA X. The sequences used for alignment in panel A are included to generate the tree. The tree is drawn to scale with branch lengths.

**S1 Table.**
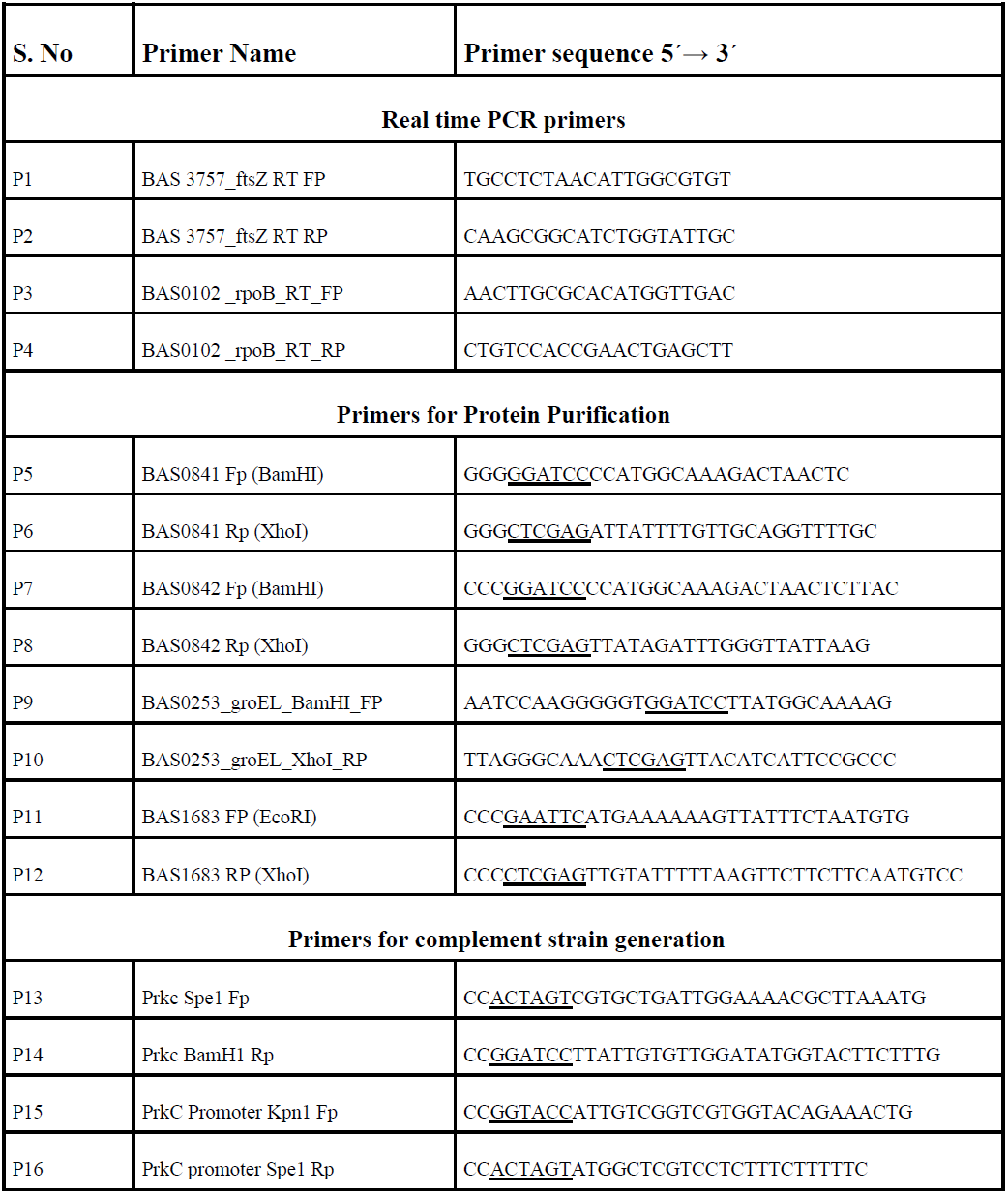
List of primers used in this study.

**S2 Table.** List of plasmids used in this study.

| S2 Table. List of plasmids used in this study |  |  |  |
| --- | --- | --- | --- |
| Name | Description | Resistance Marker | Reference |
| pYS5 | Plasmid used for complementation in *B. anthracis*; AmpR in *E. coli*; KanR in *B. anthracis* | Kanamycin, Ampicillin | [68] |
| pYS5-prkC* | Plasmid expressing PrkC under own promoter in *B. anthracis* | Ampicillin, Spectinomycin | This study |
| pProExHTc | *E. coli* expression vector with N-terminal His<sub>6</sub>-tag | Ampicillin | Invitrogen |
| pET28a | *E. coli* expression vector with N and C-terminal His<sub>6</sub>-tag | Kanamycin | Invitrogen |
| pProExHTc-sap | Expression of His<sub>6</sub>-Sap in *E. coli* | Ampicillin | This study |
| pProExHTc-eag | Expression of His<sub>6</sub>-EA1 in *E. coli* | Ampicillin | This study |
| pProExHTc-groEL | Expression of His<sub>6</sub>-GroEL in *E. coli* | Ampicillin | This study |
| pET28a-bslo | Expression of His<sub>6</sub>-BslO in *E. coli* | Kanamycin | This study |

**S3 Table.** Bacterial strains used in this study.

| S3 Table. Bacterial strains used in this study |  |  |  |
| --- | --- | --- | --- |
| Name | Genotype | Resistance marker | Reference |
| DH5α | *E. coli* F<sup>-</sup> *endA1 glnV44 thi-1 recA1 relA1 gyrA96 deoR nupG purB20* ϕ80*lacZ*Δ*M15* Δ(*lacZYA-argF*)U169, *hsdR17*(*r<sub>K</sub><sup>-</sup>m<sub>K</sub><sup>+</sup>*), λ<sup>-</sup> | - | Invitrogen |
| BL21(DE3) | *E. coli* B strain: F<sup>-</sup> *ompT gal dcm lon hsdS<sub>B</sub>*(*r<sub>B</sub><sup>-</sup>m<sub>B</sub><sup>-</sup>*) λ(DE3 [*lacI lacUV5-T7p07 ind1 sam7 nin5*]) [*malB<sup>+</sup>*]<sub>K-12</sub>(λ<sup>S</sup>) | - | Invitrogen |
| SCS110 | *E. coli* SCS110 is an *endA*<sup>-</sup> derivative of the JM110 strain *rpsL* (Strr) *thr leu endA thi-1 lacY galK galT ara tonA tsx dam dcm supE44* Δ(*lac-proAB*) [F' *traD36 proAB lacIqZΔM15*]. | - | Stratagene |
| *B. anthracis* Sterne 34F2 (BAS WT) | *B. anthracis* strain pXO1<sup>+</sup>, pXO2<sup>-</sup> | - | NIAID, NIH |
| BAS Δ*prkC* | *B. anthracis* Sterne:: *prkC* | Kanamycin | [42] |
| BAS Δ*prkC*::*prkC* | *B. anthracis* Sterne:: *prkC* + *prkC* | Kanamycin, Spectinomycin | This study |

**S4 Table.** Detailed summary statistics table for Fig 3.

| Fig 3 : Summary statistics |  |  |  |
| --- | --- | --- | --- |
| BAS WT-BAS $\Delta prkC$<br>(Hours) | 95%<br>CI of Difference | Summary | Adjusted P<br>Values |
| 2 | 79.53 to 119.3 | **** | <0.0001 |
| 3 | 69.67 to 111.1 | **** | <0.0001 |
| 4 | 69.71 to 110.4 | **** | <0.0001 |
| 5 | 28.69 to 47.28 | **** | <0.0001 |
| 6 | 14.65 to 28.93 | **** | <0.0001 |
| 7 | 14.68 to 34.50 | **** | <0.0001 |
| 8 | 8.703 to 23.49 | **** | <0.0001 |
| 9 | 7.393 to 20.47 | **** | <0.0001 |
| 10 | 0.5219 to 6.960 | * | 0.0112 |
| 14 | 14.25 to 25.90 | **** | <0.0001 |
| 18 | 12.62 to 21.36 | **** | <0.0001 |
| 22 | 5.646 to 14.16 | **** | <0.0001 |
| 26 | 2.440 to 18.08 | ** | 0.0026 |
| 30 | 1.592 to 7.368 | *** | 0.0002 |

**S5 Table.** Detailed summary statistics table for Fig 5C.

| Fig 5C : Summary statistics |  |
| --- | --- |
|  | Values |
| P Value | <0.0001 |
| Summary | **** |
| BAS WT Mean (nm) | 40.63 |
| BAS $\Delta prkC$ Mean (nm) | 33.44 |
| 95% CI | 8.101 to 6.276 |
| R squared (eta squared) | 0.5597 |

**S6 Table.** Detailed summary statistics table for Fig 6D.

| Fig 6D : Summary statistics |  |  |  |
| --- | --- | --- | --- |
| BAS WT- BAS $\Delta prkC$ | 95% CI of Difference | Summary | Adjusted P Values |
| Mid-log phase | -1.102 to 0.03703 | ns | 0.0657 |
| Stationary phase | -1.991 to -0.8526 | *** | 0.0003 |

